# Short-term hyperglycaemia induces motor defects in *C. elegans*

**DOI:** 10.1101/240473

**Authors:** Runita Shirdhankar, Nabila Sorathia, Medha Rajyadhyaksha

## Abstract

Hyperglycaemia causes various intracellular changes resulting in oxidative stress leading to loss of integrity and cell death. While cellular effects of hyperglycaemia have been reported extensively there is no clarity on whether the cellular changes translate into alterations in behaviour. Study of behavioural alterations also provides a sublime top-down approach to dapple the putative systems affected due to hyperglycaemic stress. Hence, this aspect of effect of hyperglycaemia deserves attention as it could be an early indicator of neurodegenerative changes. *Caenorhabditis elegans* is an excellent model to address these questions since it has a simple nervous system and the ability to respond to various cues.

We have investigated alteration in behaviour which involves various motor and sensory function of the *C. elegans* nervous system under hyperglycaemia. Exposure of *C. elegans* to 400 mM glucose for 4hr did not kill the worm but gave rise to decreased number of progeny, exhibiting other aberrant behaviours. This dosage was considered to cause hyperglycaemic stress and used further in the studies. Various assays that quantified behaviour, such as feeding (pharyngeal pumping/min), locomotion (distance travelled by the worms/min), olfactory response towards Butanol (response index) and gustatory response NaCl (response index) were conducted under both normal and hyperglycaemic conditions. The behavioural alterations were validated by scrutinizing changes in level of Acetylchloine which regulates motor behaviour and morphology of chemosensory neurons. Our results indicate that hyperglycaemia alters motor behaviour of the worm which was validated by a reduction in ACh levels. However, chemosensory systems were robust enough to resist reduction in neuronal integrity due to hyperglycaemic assault.

## 1. Introduction

All living creatures on earth thrive on the most indispensible compound ‘Glucose’ for their energy requirements to sustain life. Energy metabolism in mitochondria by glycolysis pathway generates major amount of ATP molecules (60–70% of total energy produced in humans (Berg, Tymoczko, & Stryer Lubert, 2002)) which are the energy currency of living systems. Life on earth will abolish without energy production by the glycolytic pathway.

However a regulated amount of this fundamental energy source needs to be maintained, failure of which has a deleterious, rather than propitious effect at the cellular levels perpetuating as system dysfunction. Low glucose concentration leads to hypoglycaemia characterized by a depressed production of energy through the glycolytic pathway making sustenance of life strenuous. On the other hand, high glucose concentration leading to hyperglycaemia is ruinous rather than favourable to living systems.

Modern day diet is mostly lopsided with high glucose intake inducing hyperglycaemia and eliciting various hyperglycaemia induced dysfunctions. An increase in sweet tooth diet due to wanton dietary habits has further contributed to an increase in hyperglycaemia in populations world-wide. Apart from these exemplifications which can be readily regulated, hyperglycaemia happens to be a manifestation of various diseases and disorders like both type I and type II diabetes, gestational diabetes, pancreatitis, Cushing's syndrome, unusual hormone-secreting tumours, pancreatic cancer, etc. Hence deciphering the mechanism that lead to cellular assault resulting in system dysfunction in humans becomes crucial to design drugs and therapies for rescue from hyperglycaemic stress.

### 1.1. Pathology of hyperglycaemic stress

Incursion by hyperglycaemia results into various cellular changes before it finally activates the programmed cell death pathway. It causes an increase in superoxide formation, increasing the oxidative stress due to increased formation of ROS species. This increase in ROS species further makes most of the cellular pathways chaotic eventually compelling the cell to undergo apoptosis (Dobretsov, Romanovsky, & Stimers, 2007).

Hyperglycaemia activates a number of enzymatic and non-enzymatic pathways of glucose metabolism that include

1. Increased polyol pathway activity leading to sorbitol and fructose accumulation, NAD(P)-redox imbalances, and changes in signal transduction
2. Non-enzymatic glycation of proteins yielding “advanced glycation end-products” (AGEs);
3. Activation of protein kinase C (PKC), initiating a cascade of stress responses
4. Increased hexosamine pathway flux
5. Hyperglycaemia-mediated superoxide overproduction by the mitochondrial electron transfer chain. Each of these mechanisms result in reactive oxygen species (ROS), reflected in an overall increased state of cellular oxidative stress (Vincent, McLean, Backus, & Feldman, 2005).

Mitochondria when observed under electron microscope appear swollen with its inner cristae disrupted, hence, making it evident that the production of ROS species in the ETC cycle due to increased glycolytic flux induces the increased oxidative stress in the cell during hyperglycaemic assault (Sivitz & Yorek, 2010).

Various combat mechanisms reinforced to rescue the cell from the increased oxidative stress. With an increase in ROS production the production of antioxidants in the cell is upregulated. The administration of traditional antioxidants, vitamins A, C, and E and alpha lipoic acid, that have the capacity to rapidly scavenge a variety of free radicals (Oyenihi, Ayeleso, Mukwevho, & Masola, 2015). Glutathione level goes up to reduce the formation of advanced glycation end products by the glyoxalase system that converts the methyl glyoxal formed into lactate (Vincent et al., 2005). There is an increase in enzymes such as superoxide dismutase, catalase, etc. which aid regulating the ROS levels in the body (Vincent et al., 2005). However, with at higher glucose concentrations of hyperglycaemic conditions for a prolonged period of time these rescue mechanisms eventually fall apart which fatal to the cell.

### 1.2. Hyperglycaemia induced neuropathy

Since neurons barely divide or regenerate post mitotically, with the exception of a few privileged neuron types in the hippocampus, olfactory bulb, etc., a loss of these non-dividing neurons carrying out crucial and paramount functions of the body would be debilitating.

The ever bustling neurons of the human brain have the highest energy demand which require a continuous delivery of glucose from blood. The human brain even though accounts for as small as ~2% of the body weight, consumes almost ~20% of glucose-derived energy making it the major consumer of glucose. Hence, tight regulation of glucose metabolism is critical for brain physiology and disturbed glucose metabolism in the brain underlies several diseases affecting both the brain itself as well as the entire organism (Mergenthaler, Lindauer, Dienel, & Meisel, 2013).

Hyperglycaemic stress induces various distressful alterations in cellular mechanism of neurons which reduce neuronal integrity affecting the functioning of organism as a whole. Even more catastrophic is apoptosis of neurons in fatal hyperglycaemic conditions wherein, the neurodegeneration of neurons adversely and irreversibly affects the function carried out by the specific neurons. Hence, the mechanism of hyperglycaemia induced neuropathy needs to be deciphered to device methods for its prevention.

### 1.3. Behaviour to study hyperglycaemic neuropathy

Different types of neurons throughout the brain exhibit varying degree of robustness on predisposition to stress conditions. Even subtle differences in characteristics of neurons like various receptors, cell membrane composition, neurotransmitters involved, etc., results in difference in susceptibility to damage due to stress conditions. Damage to specific type of neurons would lead to its manifestation as an aberrant behaviour which is associated with that particular neuron type.

The intricacy of the nervous system made of millions of neurons forming trillions of synapses makes it difficult to get a fix on the specific neurons affected due to any stress condition. An easier path to recognize the affected neurons is a top-down approach to first study various different behaviours for abnormal patterns to identify the putative neurons associated with such aberrant behaviours and further conduct a detailed study of the cellular and molecular alterations occurring in these potentially affected neurons only.

### 1.4. Hyperglycaemia in *C. elegans*

**Fig. 1.**
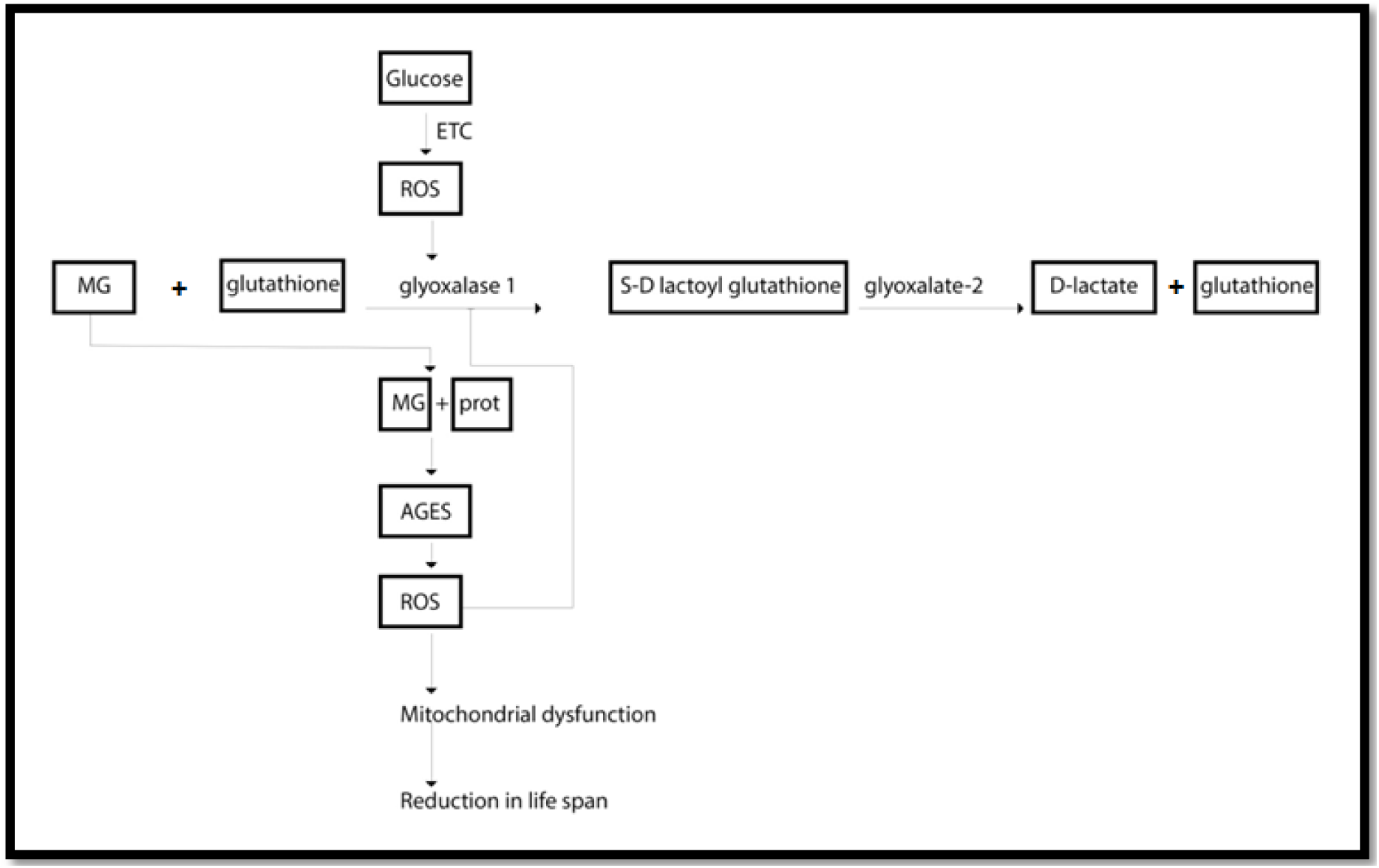
Glyoxalase system dysfunction induced due to Hyperglycaemic assault in mitochondria Image designed using software Corel Draw Graphics Suite X5.

The effect of hyperglycaemia on the glyoxalase system has been well studies using model organism *C. elegans*. In *C. elegans* increased glycolytic flux in hyperglycaemic conditions leads to formation of mitochondrial reactive oxygen species (ROS). In turn, ROS formation decreases the activity of glyoxalase-1, an enzyme detoxifying methylglyoxal. Accumulation of methylglyoxal, a highly reactive dicarbonyl derived from triosephosphates, leads to rapid protein modification. Methylglyoxal is an arginine-directed glycating agent and precursor of advanced glycation end products (AGEs). It modifies proteins mainly but not exclusively on arginine residues forming methylglyoxal-derived hydroimidazolone (MG-H1) with formation of other minor AGE residues, arginine-derived ‘ argpyrimidine and lysine-derived N-(1carboxyethyl)lysine and methylglyoxal-derived lysine dimer. Methylglyoxal-derived modification of mitochondrial proteins increases mitochondrial ROS formation, thus reducing life span in *C. elegans*. The formation of methylglyoxal-modified proteins in cells is suppressed by the activity of the enzyme glyoxalase-1 that catalyzes conversion of methylglyoxal, with the cofactor glutathione, to S-D-lactoylglutathione. This is further converted to d-lactate by glyoxalase-2 regenerating the glutathione (Schlotterer et al., 2009).

### 1.5. Behaviours in *C. elegans* to study hyperglycaemia

*C. elegans* is a free living soil nematode with a primitive and simple body structure and functions and yet shares relatively substantial amount of homology with humans. Its simple nervous system made of computable amount of neurons makes it an exquisite model organism for neurobiological studies. 302 neurons although might sound like a number easy to investigate, takes an accountable amount of labour, time and finances. Hence to narrow down the effect of hyperglycaemia on the nervous system to only the putative neurons the top-down approach of analysing the different simple behaviours exhibited by *C. elegans* is the most favourable approach. *C. elegans* exhibits various simple behaviours (Hart, 2006).

#### Motor behaviour

The motor system of *C. elegans* exhibits simple motor behaviours required for moving around, feeding and reproduction. The locomotion of the worms is characterized by an elegant (hence the name) sinusoidal motion by alternating ventral and dorsal musculature turns. Feeding in *C. elegans* is also controlled by the activity of motor neurons that regulate the rate of pharyngeal pumping and isthmus peristalsis for food ingestion. The vulva and uterine muscles control egg laying in worms which is regulated by motor neuron (HSN and VC).

Locomotion: The neuromuscular system controlling locomotion creates a pattern of contraction and relaxation. Locomotion is based on sensory inputs. A typical body movement of *C. elegans* is characterized by a rhythmic pattern and cyclic waves. Environmental cues regulate wave frequency, amplitude, wavelength and rate of propagation (Leifer, Butler, Fang-yen, Kawano, & Samuel, 2013). The ventral nerve cord subcircuit associated with forward locomotion contains four main classes of motor neurons: 11 neurons of class VB, 7 DB, 13 VD, and 6 DD as well as two key pairs of command interneurons (classes AVB and PVC) (Cohen N. et.al. 2014).

Feeding: Feeding behaviour is controlled by pharynx, a neuromuscular pump that transports bacteria from moth of the worm to its intestine. It consists of two types of motion the terminal bulb pump and the isthmus peristalsis. Pharyngeal pumping is a cycle of contractions and relaxation caused by a single action potential responsible for sucking in food and water from surrounding environment done by pharynx which is controlled by three types of neurons located below the basal lamina, two (MC and M3) presiding over pharyngeal pumping and one (M4) over isthmus peristalsis (Song & Avery, 2013). MC neuron induces contraction of corpus and this waves travels to the terminal bulb initiating its contraction. The I1 neuron fine tunes the firing of the MC neuron.

#### Chemosensation

*C. elegans* detect miniscule concentrations of volatile odorants using olfactory system and soluble compounds using gustatory system which is linked to detecting food, avoid noxious conditions, develop appropriately, mate, etc. 5% of the total genes are involved in chemosensation with substantial number of neurons involved. Sensation of these chemicals influences chemotaxis, rapid avoidance, overall change in motility and even entry or exit from dauer stage. Each amphid sensory neuron expresses a specific set of candidate receptor genes that detect attractants, repellents and pheromones

Gustation: Gustatory system is responsive to chemical agents like cations (Na^+^), anion (Cl^−^), cyclic nucleotides (cAMP) and amino acids (serotonine). ASE (Amphid Sensory) neurons are the primary neurons involved in sensing tastes of chemotactic agents. Other than ASE neuron ADF, ASG, ASI, ASK, ASJ, etc. are also involved in gustation.

Olfaction: AWA, AWB and AWC sensory neurons are involved in detection of volatile odors in concentrations as low as nanomolar hence exhibits long range chemotaxis. *C. elegans* can smell alcohol,, ketones, aldehydes, esters, amines, sulfydryls, organic acids, aromatic and hetrocyclic compounds which are all present as natural products of bacterial metabolism.

### 1.6. Role of acetylcholine in motor system

*C. elegans* uses acetylcholine at neuromuscular synapses to control muscle contraction. In a wild-type animal, acetylcholine released from cholinergic motor neurons stimulates postsynaptic cholinergic receptors of the body-wall muscles causing them to contract. Coordinated exocytosis of acetylcholine allows *C. elegans* to move nearly continuously in a sinusoidal manner. The pharynx generates a myogenic rhythm in the presence of tonically released acetylcholine. The MC neurons that control the pharyngeal pumping appear to act primarily via a nicotinic acetylcholine (ACh) receptor containing the non-α subunit EAT-2 (Trojanowski, Raizen, & Fang-Yen, 2016).

## 2. Methods and Materials

### i. Maintenance and synchronization of worm cultures

<u>Maintenance:</u> N2 Bristol strain of *C. elegans* was used. The worms were cultured on standard Nematode Growth Medium (17g agar, 3g NaCl, 2.5g peptone, 1ml 1MCaCl_2_, 1ml 1M MgSO_4_, 25ml 1M KPO_4_ buffer, dilute with water up to 1litre (Zhu et al., 2016)) at 22°C seeded with *Escherichia coli* OP50 (Stiernagle, 2006). Worms were subcultured by the washing technique once every 3 days wherein, sterile M9 buffer (3g KH_2_PO_4_, 6g Na_2_HPO_4_, 5g NaCl, 1ml 1M MgSO_4_, dilute with water up to 1litre (Zhu et al., 2016)) was used to extract the worms out of the plate. The M9 buffer containing worms was transferred to a test tube, a pellet containing the worms was allowed to form, repeated washed of sterile M9 buffer were given and the pellet containing the worms was finally transferred to a fresh NGM plate.

**Fig.2.**
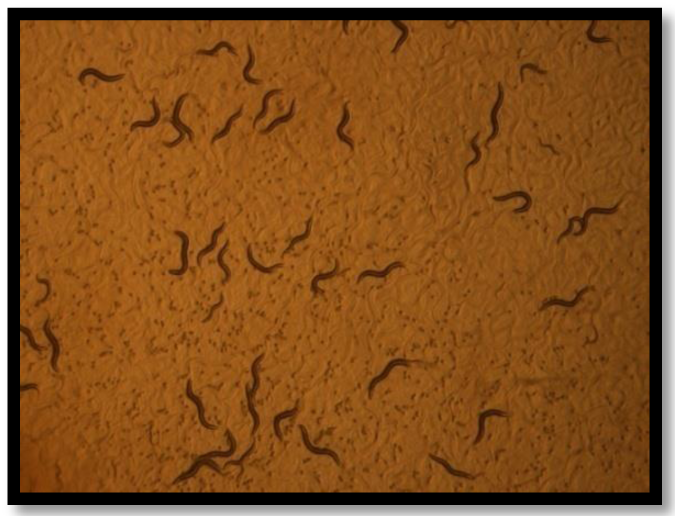
NGM plate containing adult worms and eggs Image captured using NIKON SMZ1500 stereodissection microscope at magnification 5x.

<u>Synchronization:</u> For synchronization of worm culture, egg-laying adult *C. elegans* were washed with sterile M9 buffer and the worms collected were transferred to an eppendorf. 0.2ml of 5N NaOH and 0.4ml of 5% sodium hypochlorite was added and diluted up to 1ml using M9 buffer. The mixture was centrifuged at 4000rpm for 3 minutes to form a pellet with released eggs. The supernatant was aspirated out to 0.1ml. Eggs in the remaining 0.1ml of liquid were repeatedly washed with M9 buffer before it was transferred to NGM plate pre seeded with *E.coli* OP50.

### ii. Glucose standardization assay

Various concentrations of glucose (0mM, 100mM, 200mM, 300mM, 400mM and 500mM) were used and the viability of worms was checked at different time points (0hr, 2hr, 4hr and 24hr).

All assays were carried out using synchronised worms on first day of adult cycle.

Untreated worms were ones without glucose exposure.

Treated worms were exposed to 400mM of glucose (G8270 SIGMA-ALDRICH) for 4 hours. Glucose was prepared using K medium (Zhu et al., 2016) (2.2365g KCl, 2.922g NaCl, 0.8405g NaOAc, dilute with water up to 1litre) and introduced to NGM plate using spread plate technique and pre-seeded with *E. coli* OP50.

### iii. Behavioural assay

#### a. Motor behaviour assay

##### I. **Locomotion assay** (Hart, 2006)

Single worms, both untreated and treated, were transferred to 6cm assay plates containing NGM and their body movement was recorded using NIKON SMZ1500 stereodissection microscope at 5x and ACT-2U software.

The distance travelled by the worms were measures using the software ImageJ.

##### II. **Pharyngeal Pumping assay** (Trojanowski et al., 2016)

Single worms, both untreated and treated, were transferred to 6cm assay plates containing NGM and their pharyngeal pumping was recorded using NIKON SMZ1500 stereodissection microscope at 20x and ACT-2U software.

The rate of pharyngeal pumping was measured manually.

#### b. Chemosensory behavior assay

##### I. **Olfactory assay** (Margie, Palmer, & Chin-Sang, 2013)

Assay was carried out using both untreated and treated worms. The underside of 6cm assay plate was divided into 4 equal quadrants and a circle with radius 0.5cm at the center. Two diagonally opposite quadrants were marked as ‘T’ (test) and the remaining two as ‘C’ (control) for test plates and all quadrants as ‘C’ (control) for control plates close to the edge of the plate at the center of the curvature. Chemotaxis agar (1.7%agar, 5mM KPO_4_ buffer, 1mM CaCl_2_, 1mM MgSO_4_) was used. The test solution was prepared by mixing 1% ‘Butan-1-ol’ (SD fine chemicals limited) with equal volume of 0.5 M sodium azide (an anesthetic used to arrest the worms upon reaching a quadrant). The control solution was prepared by using equal volume of distilled water and 0.5 M sodium azide. 10μl of the worm was loaded onto the center of the plate within the marked circle. Immediately after, 2μl of the control solution was introduced onto the two control sites and all four control sites of the control plates. Likewise, the same amount of the test solution was introduced onto the two test sites. The plates were covered with the lids immediately and left in the dark at 22°C for an hour. The number of worms in each quadrant was counted after an hour under NIKON SMZ1500 stereodissection microscope at 5x.

Interpreting the Scores/Determining the Chemotaxis Index: The chemotaxis response index was calculated.

##### II. **Gustatory assay** (Hart, 2006)

Assay was carried out using both untreated and treated worms. Four quadrant assay plates were used. Two diagonally opposite quadrants were marked as ‘T’ (test) and the remaining two as ‘C’ (control) for test plates and all quadrants as ‘C’ (control) for control plates. Chemotaxis agar (1.7%agar, 5mM KPO_4_ buffer, 1mM CaCl_2_, 1mM MgSO_4_) was used in control quadrants and chemotaxis agar with 50mM NaCl was used in test quadrants. 10μl of the worms were loaded onto two of the linear partitioning between the test and control quadrants. The plates were covered with the lids immediately, and left in the dark at 22°C for one hours. The number of worms in each quadrant was recorded after two hours under NIKON SMZ1500 stereodissection microscope at 5x.

Interpreting the Scores/Determining the Chemotaxis Index:

The chemotaxis response index was calculated.

##### iv. **Estimation of specific activity of Acetylcholine esterase by Ellman’s method** (Ellman, Courtney, Andres, & Featherstone, 1961)

Protein extraction: 100μl of worm pellet was used for both treated and untreated worms. The pellet was macerated using 1ml of 0.1M phosphate buffer pH 8.0. This mixture was centrifuged at 15,000 r.p.m at 4°C for 15 minutes. The supernatant was used to estimate protein by The Folin Lowry method of protein estimation.

Enzyme estimation: To 2.6ml of 0.1M phosphate buffer pH 8.0, 400μl of the supernatant was added. 100μl DTNB was added and the absorbance at 412nm was measured and set to zero. 20μl of Acetylthiocholine Iodide substrate was added. Absorbance was recorded at 412nm every 10minutes. Specific activity of the enzyme acetylcholine esterase was calculated for both test and control.

##### v. Statistical Analysis

Statistical analysis was performed by applying a two-tailed Student’s t-test to compare the high-glucose treated group and the non-treated group. Software Excel 2010 was used in this work.

## 6. Observations and Results

### i. Glucose standardization assay

An increase in both dose and time of exposure to glucose causes an increase in the rate of mortality in *C. elegans*.

**Mortality Rate at Different Time Points:** An increase in both dose and time of exposure to glucose causes an increase in the rate of mortality in *C. elegans*.

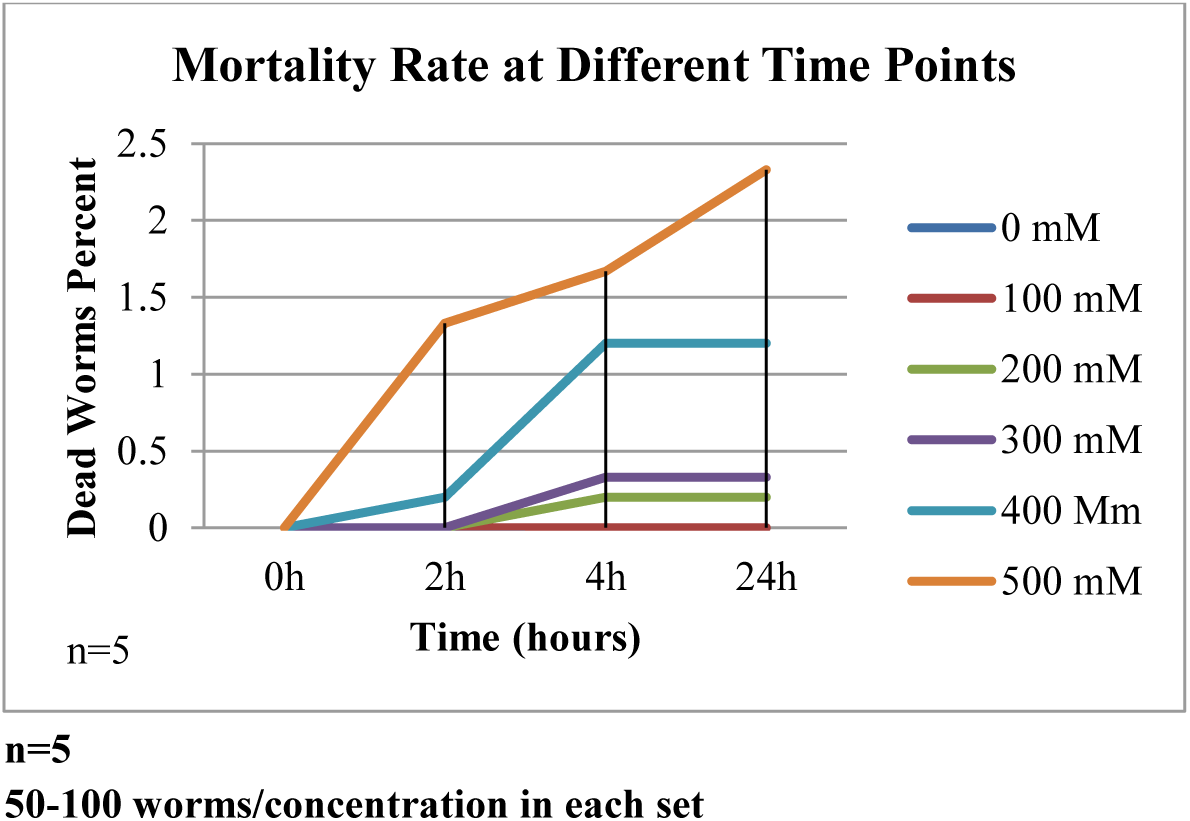

**Mortality Rate at 4^th^ hour:** A significant decrease in the rate of mortality was observed in worms exposed to glucose concentratiosn above 200mM concentration of glucose at the 4^th^ hour. The values represent mean +/− SE. Students t test for unpaired values: **p<0.01 and ***p< 0.001 between worms without glucose exposure and worms exposed to various doses of glucose.

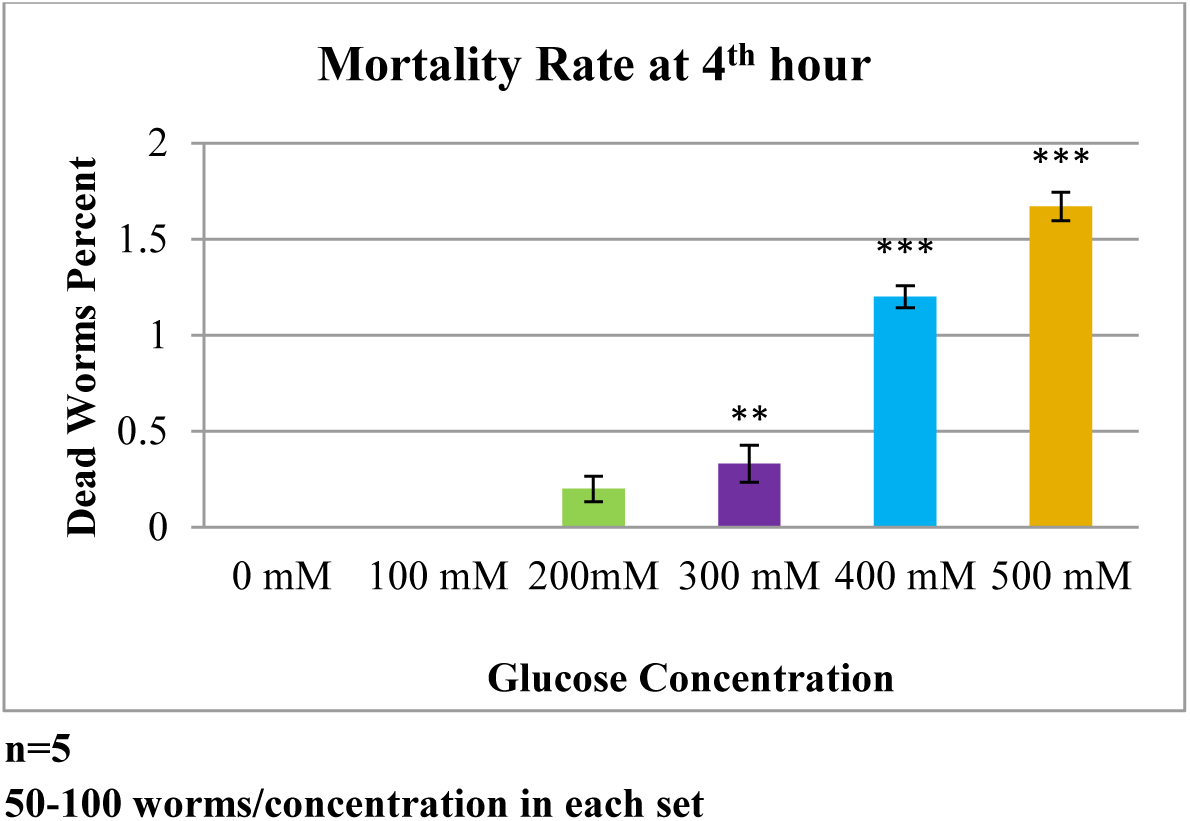

**Aberrant behaviour:** Curling behaviour which is a characteristic aberrant behaviour observed in *C. elegans* under stress conditions was observed susbtantially at 400mM concentration of glucose.

**Fig.3.**
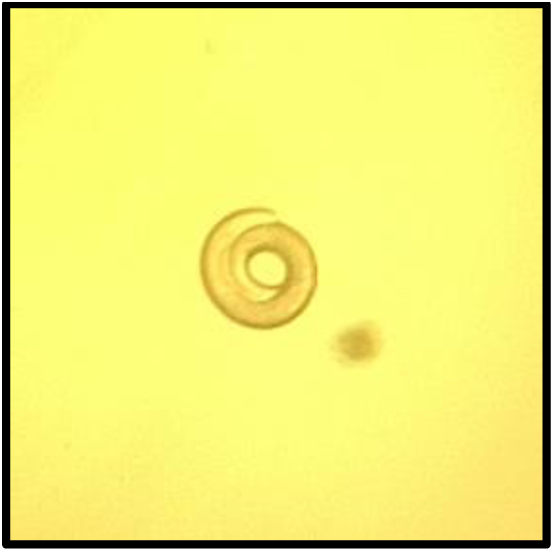
Curling behavior observed at 400mM Image captured using NIKON SMZ1500 sterodissection microscope and software ACT-2U at 5x

### ii. Behavioural assay

#### a. Motor behaviour assay

**I. Locomotion assay:** A significant decrease in locomotion behavior was observed in hyperglycemic worms. The values represent mean +/− SE. Students t test for unpaired values: ***p< 0.001 between untreated and treated worms.

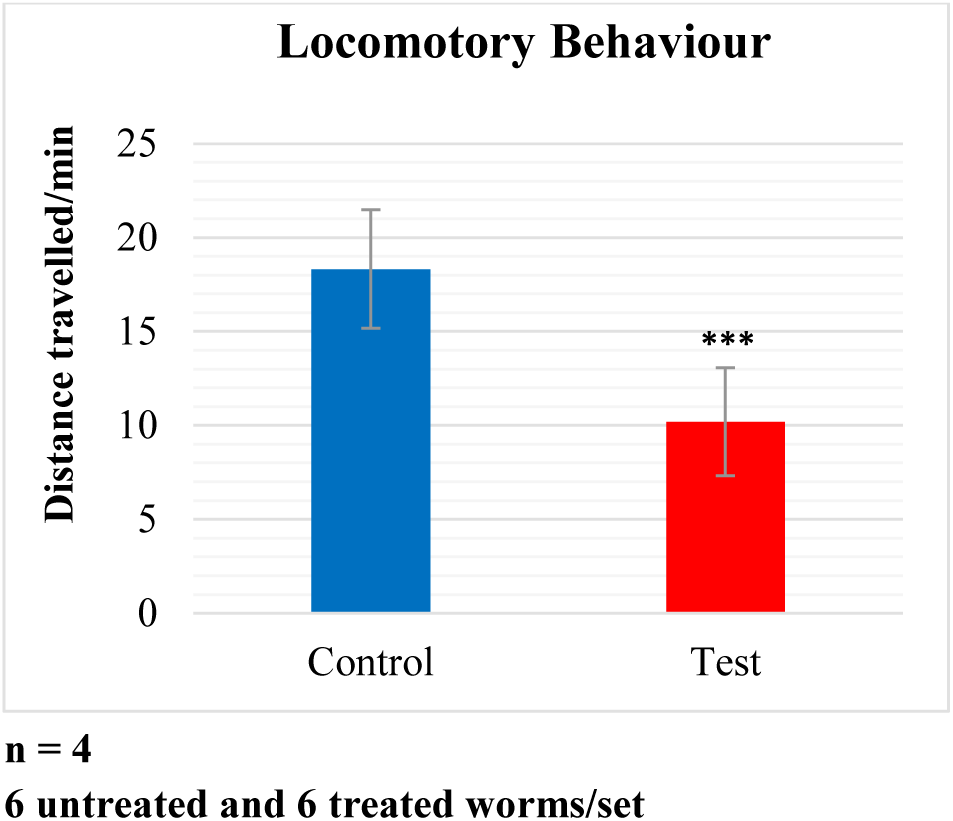

**II. Feeding assay:** A significant decrease in pharyngeal pumping was observed in hyperglycemic worms. The values represent mean +/− SE. Students t test for unpaired values: ***p< 0.001 between untreated and treated worms.

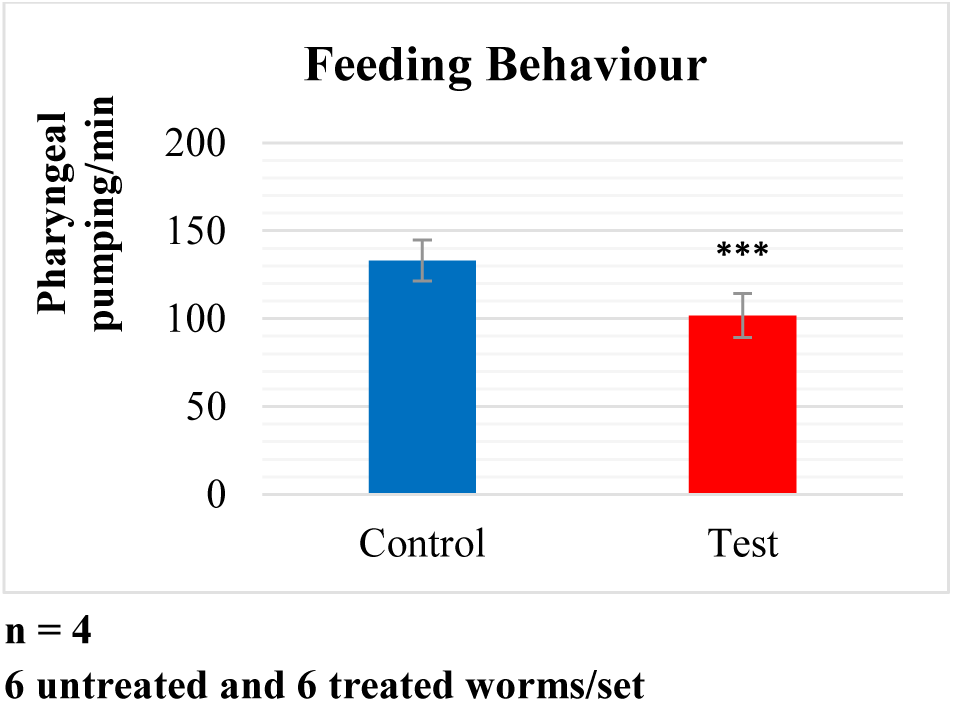

#### b. Chemosensosry behaviour assay

**I. Olfactory assay:** No significant difference was observed in chemosensory response towards Butanol. The values represent mean +/− SE.

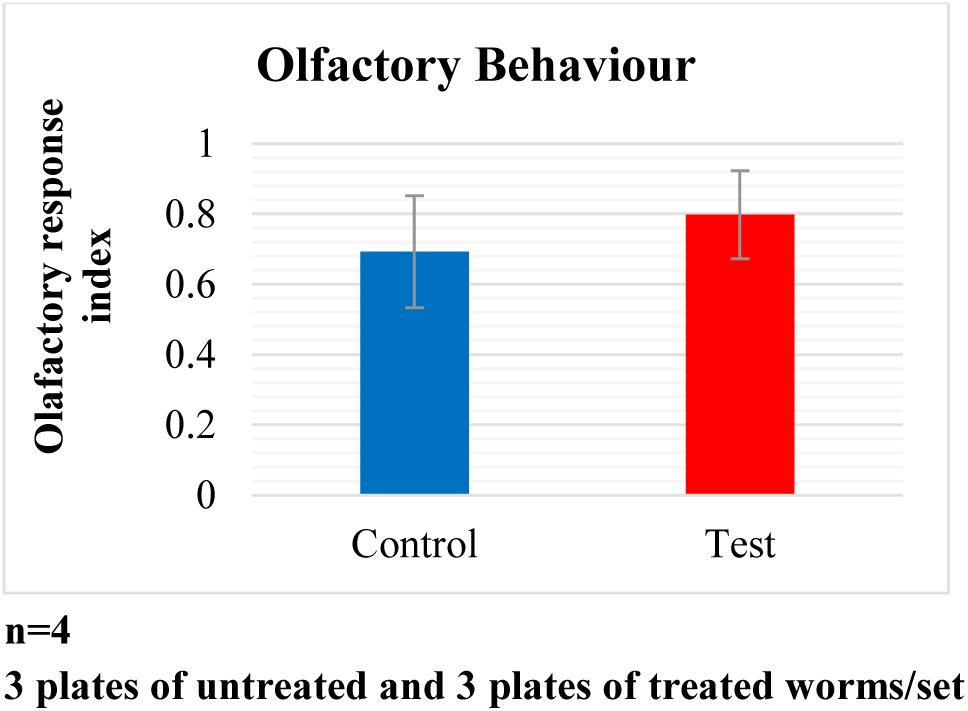

**II. Gustatory assay:** No significant difference was observed in chemosensory response towards NaCl. The values represent mean +/− SE.

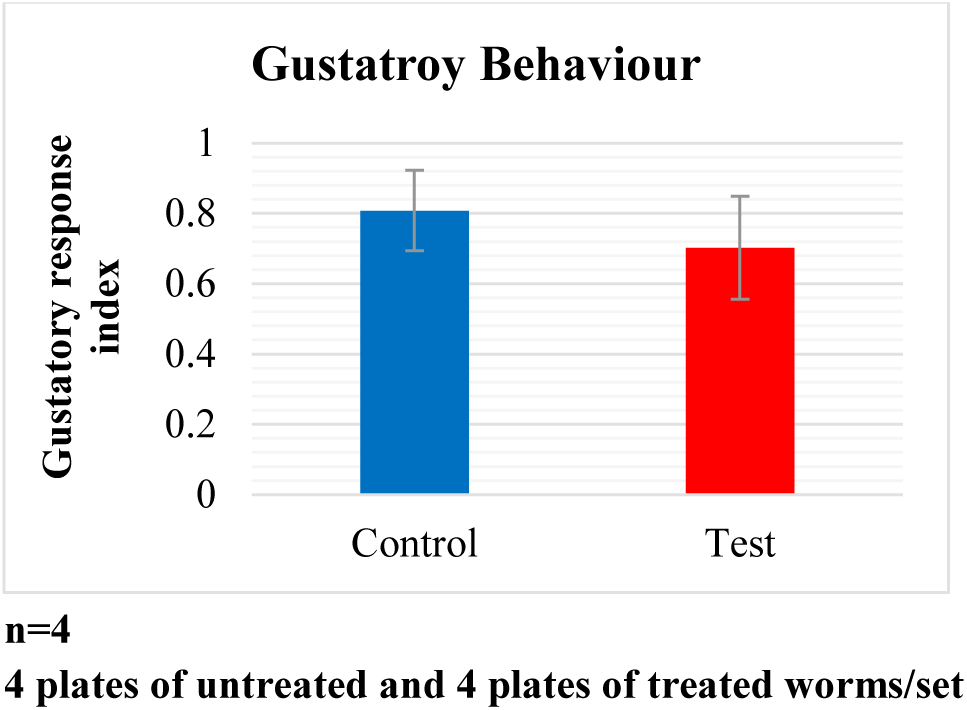

**iii. Estimation of acetylcholine esterase activity:** A significant decrease in specific activity of enzyme acetylcholine esterase was observed in hyperglycemic worms. The values represent mean +/− SE. Students t test for unpaired values: *p< 0.05 between untreated and treated worms.

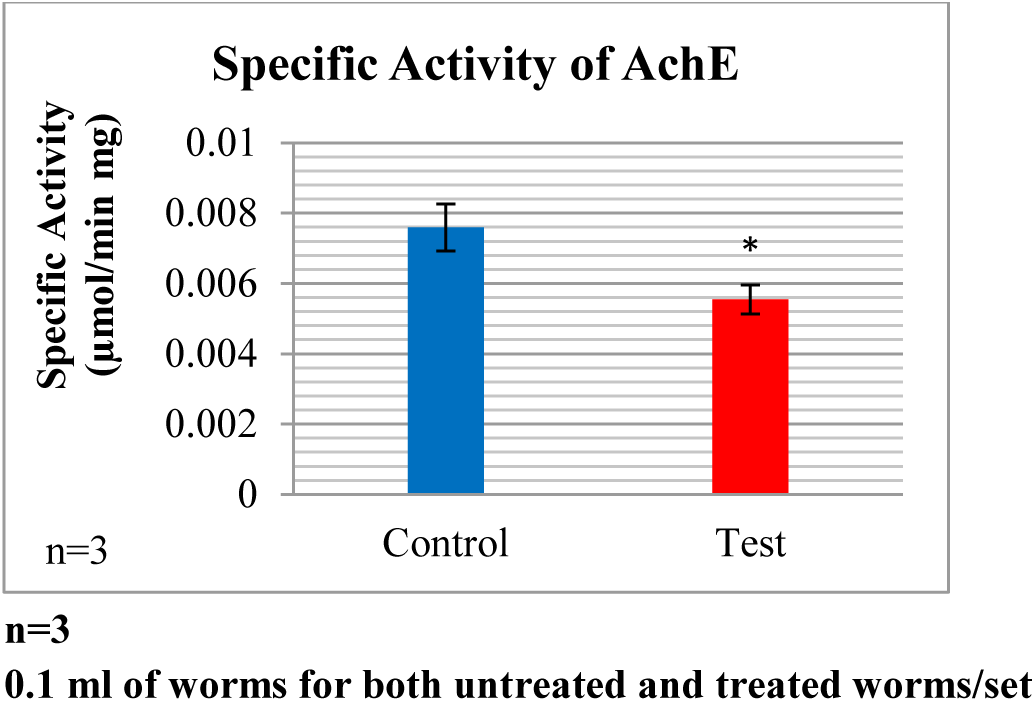

## 7. Discussions

### i. Standardization of glucose dosage

Mortality at 100mM was absent. At 200mM and 300mM very little mortality rate was observed at the 4^th^ hour which remained stable up to the next 24 hours. At 400mM and 500mM was observed first at the 2^nd^ hour itself. For 400mM mortality rate increased till the 4^th^ hour and was unchanged until the 24^th^ hour. However, for 500mM the rate went on increasing even until the 24^th^ hour proving to be a fatal dose of glucose. Hence 4 hour exposure time was chosen as the standard time of exposure since the rate of mortality remained constant after the 4^th^ hour up to 400mM dose of glucose.

At the 4^th^ hour mortality rate was significantly different from mortality rate without glucose treatment and also remained stable until the 24^th^ hour. Hence 400mM of glucose was chosen as the standard dose of glucose.

### ii. Effect of hyperglycaemia on Motor System

Both locomotion and pharyngeal pumping which are activities controlled majorly by the motor system were affected due to hyperglycaemia implying that hyperglycaemic assault leads to degeneration of the neuromuscular system.

To further investigate and validate the degeneration of neuromuscular system the activity of acetylcholine was indirectly checked by investigating the specific activity of acetylcholine esterase, the activity of which positively corresponds to the activity of acetylcholine, the major neurotransmitter involved in motor function in *C. elegans*. a decrease in the specific activity of acetylcholine esterase was found, implying that hyperglycaemic stress induces decreased acetylcholine activity.

### iii. Effect of hyperglycaemia on Sensory System

Both chemosensory behaviours, gustatory and olfactory, were found to be unaffected by hyperglycaemia treatment signifying that the chemosensory system is robust and has the ability to combat hyperglycaemic assault.

## 8. Conclusion

Short-term hyperglycaemia leads to neuropathy of the motor system which could possibly be due to reduced activity of major neurotransmitter acetylcholine. However, the chemosensory system is robust against hyperglycaemic stress and morphological changes in the chemosensory neurons are to be investigated further.

## 9. Acknowledgement

We would like to acknowledge the support of DST-FIST funds for infrastructure and DBT-Neuroscience funds for consumables are acknowledged. And Dr. Momna Hejmadi, University of Bath, U.K for providing us with the worm cultures.

## References

• Berg, J. M., Tymoczko, J. L., & Stryer Lubert. (2002). Biochemistry. (W. H. Freeman, Ed.) (5th Editio). New York. Retrieved from http://www.whfreeman.com/

• Dobretsov, M., Romanovsky, D., & Stimers, J. R. (2007). Early diabetic neuropathy: Triggers and mechanisms. World Journal of Gastroenterology, 13(2), 175–191. https://doi.org/10.3748/wjg.v13.i2.175

• Ellman, G. L., Courtney, K. D., Andres, V., & Featherstone, R. M. (1961). A new and rapid colorimetric determination of acetylcholinesterase activity. Biochemical Pharmacology, 7, 88–95. https://doi.org/10.1016/0006-2952(61)90145-9

• Hart, A. (2006). Behavior. WormBook, 1–67. https://doi.org/10.1895/wormbook.1.87.1

• Leifer, A. M., Butler, V., Fang-yen, C., Kawano, T., & Samuel, A. D. T. (2013). Proprioceptive coupling within motor neurons drives C. elegans forward locomotion. Neuron, 76(4), 750–761. https://doi.org/10.1016/j.neuron.2012.08.039.Proprioceptive

• Margie, O., Palmer, C., & Chin-Sang, I. (2013). *C. elegans* Chemotaxis Assay. Journal of Visualized Experiments, (74), 1–6. https://doi.org/10.3791/50069

• Mergenthaler, P., Lindauer, U., Dienel, G., & Meisel, A. (2013). Sugar for the brain: the role of glucose in physiological and pathological brain function. Trends in Neuroscience, 36(10), 587–597. https://doi.org/10.1016/j.tins.2013.07.001.Sugar

• Oyenihi, A. B., Ayeleso, A. O., Mukwevho, E., & Masola, B. (2015). Antioxidant strategies in the management of diabetic neuropathy. BioMed Research International, 2015. https://doi.org/10.1155/2015/515042

• Schlotterer, A., Kukudov, G., Bozorgmehr, F., Hutter, H., Du, X., Oikonomou, D.,… Morcos, M. (2009). C. elegans as Model for the Study of High Glucose-Mediated Life Span Reduction. Diabetes, 58(November). https://doi.Org/10.2337/db09-0567.A.S.

• Sivitz, W. I., & Yorek, M. A. (2010). Mitochondrial dysfunction in diabetes: from molecular mechanisms to functional significance and therapeutic opportunities. Antioxidants & Redox Signaling, 12(4), 537–77. https://doi.org/10.1089/ars.2009.2531

• Cohen, N., & Sanders, T. (2014). Nematode locomotion: Dissecting the neuronal-environmental loop. Current Opinion in Neurobiology, 25, 99–106. https://doi.org/10.1016Zj.conb.2013.12.003

• Song, B., & Avery, L. (2013). The pharynx of the nematode *C. elegans:* A model system for the study of motor control. Worm, 2(1), e21833. https://doi.org/10.4161/worm.21833

• Stiernagle, T. (2006). Maintenance of C. elegans. WormBook: The Online Review of C. Elegans Biology, (1999), 1–11. https://doi.org/10.1895/wormbook.1.101.1

• Trojanowski, N. F., Raizen, D. M., & Fang-Yen, C. (2016). Pharyngeal pumping in Caenorhabditis elegans depends on tonic and phasic signaling from the nervous system. Scientific Reports, 6(February), 22940. https://doi.org/10.1038/srep22940

• Vincent, A. M., McLean, L. L., Backus, C., & Feldman, E. L. (2005). Short-term hyperglycemia produces oxidative damage and apoptosis in neurons. FASEB Journal: Official Publication of the Federation of American Societies for Experimental Biology, 19(6), 638–40. https://doi.org/10.1096/fj.04-2513fje

• Zhu, G., Yin, F., Wang, L., Wei, W., Jiang, L., & Qin, J. (2016). Modeling type 2 diabetes-like hyperglycemia in C. elegans on a microdevice. Integrative Biology, 8, 30–38. https://doi.org/10.1039/c5ib00243e

